# Barrier Function of the Extracellular Matrix in AAV Gene Therapy

**DOI:** 10.1101/2025.08.01.668133

**Authors:** Yahya Cheema, Devorah Cahn, Sahana Kumar, Matthew Wolf, Gregg A. Duncan

## Abstract

**Purpose:** The extracellular matrix (ECM) is a major component of the tissue microenvironment which may pose a barrier to the distribution of AAV in target organs, preventing delivery of therapeutic cargo. We sought to address this potential barrier to AAV gene therapy by furthering our understanding of AAV-ECM interactions. We hypothesized that both the AAV serotype and ECM composition will impact AAV transport and gene delivery.

**Methods:** AAV2, AAV6, and AAV8 viral vectors were fluorescently labeled to allow for visualization of their diffusion through the ECM. Lung, liver, and small intestinal submucosal dECM hydrogels were formulated as models of the ECM with tissue-specific biomolecular content. We then characterized AAV and nanoparticle diffusion within decellularized ECM using fluorescent video microscopy and multiple particle tracking. Additionally, we evaluated AAV transduction in dECM-incorporated 2D and 3D spheroid tissue culture models.

**Results:** All AAV displayed reduced diffusivity through ECM as compared to similarly sized nanoparticles. AAV2 diffusion was least affected by the presence of ECM across tissue types as compared to AAV6 and AAV8. AAV transduction in dECM incorporated *in vitro* models was significantly reduced in both a 2D and 3D setting.

**Conclusions:** These results suggest binding of AAV to the ECM may decrease their therapeutic effect in target tissues throughout the body. The barrier function of the ECM should be considered in development of AAV for gene therapy applications.

## Introduction

Adeno-associated virus (AAV) has emerged as a leading viral gene delivery system due to the large number of available serotypes that are highly effective at transducing (infecting) diverse tissue types (1,2). AAV vectors have been employed in several clinical trials targeting conditions including cystic fibrosis, hemophilia, and muscular dystrophy (3). Given its broad potential use, it is important to understand the distribution of AAV serotypes following systemic or local injection as this will influence the efficiency of gene delivery and ultimately, its therapeutic effect (4,5). Once AAV vectors exit the bloodstream and extravasate into target tissues, the extracellular matrix (ECM) microenvironment contains many biological polymers including collagen, fibronectin, laminin, and various proteoglycan species, that presents both a physical and adhesive barrier to AAV viral particle transport to target cells. It is expected adhesive trapping of AAV within the tissue interstitial spaces to be mediated by counter-receptors, such as heparan sulfate and sialic acid, found abundantly in proteoglycans within the ECM (6,7). Moreover, past work has sought to harness ECM biomaterials for localized gene delivery through non-covalent binding and incorporation of AAV in ECM scaffolds composed of collagen, laminin, and elastin (8–11). However, the extent to which the ECM presents a barrier to gene delivery for specific AAV serotypes has yet to be clearly established.

There is compelling evidence from past studies that the ECM is an important biological barrier to consider in development of AAV as a gene therapy platform. For example, prior work has illustrated AAV transduction can be enhanced in the eye when these vectors are co-infused with ECM-degrading enzymes (e.g. collagenase, hyaluronidase) (12). Moreover, the distribution of AAV transduction within the brain, a tissue highly enriched in ECM, is significantly hindered as compared to similarly sized synthetic nanoparticles following convective delivery to the cortical regions of the brain (13). While this demonstrates that the ECM likely inhibits AAV transduction, it remains unclear to what extent ECM barrier function is generalizable across all tissues. This uncertainty arises because the composition and organization of the ECM within interstitial spaces is known to be highly tissue specific (14). Furthermore, disease-associated changes to the ECM, such as increased deposition or remodeling, may further augment its barrier function towards AAV vectors. For example during the progression of liver disease, the composition and abundance of ECM in the liver change significantly as the tissue becomes fibrotic with formation of scar tissue consisting of fibrillar collagens, fibronectin, proteoglycans, and basement membrane components (15,16). In recent work, this has been implicated in the observed reductions in hepatic gene delivery in mouse models of liver fibrosis where AAV transgene expression was inversely correlated with the extent of collagen deposition (17). However, from an *in vivo* context, it is difficult to isolate the impact of tissue specific ECM properties on AAV transduction and trafficking within target organs.

To determine the effect of ECM as an extracellular barrier to AAV gene delivery, we assessed the diffusive and transduction capabilities of multiple AAV serotypes in decellularized ECM (dECM) incorporated *in vitro* models. We selected 3 AAV serotypes, AAV2, AAV6, and AAV8 to evaluate in these studies as all are currently in preclinical and clinical development for gene therapy applications (3,18). It should also be noted each of these serotypes possess distinct binding affinities for receptors which have a strong influence on their tissue tropism. Tissue tropism can be defined as the propensity of an AAV serotype to infect specific tissue types following systemic injection. AAV2 has a strong preference for heparan sulfate whereas AAV6 binds to both heparan sulfate and sialic acid containing proteoglycans to mediate cell entry (6,7). Alternatively, AAV8 does not utilize either heparan sulfate or sialic acid for cell entry with its primary receptor being the 37/67-kDa laminin receptor (LamR) (19). dECM hydrogels were also used as a scaffold material such that our models included the full complement of biomolecular components found within the ECM of native tissues (20). This study provides direct evidence that AAV can become trapped within ECM from various tissues, resulting in diminished transduction efficiency. The results of these studies provide the basis for further advancements in capsid engineering and vector administration to circumvent the ECM barrier to AAV gene therapy.

## Materials and methods

### Decellularized Extracellular Matrix Hydrogel Formation

Porcine lung, liver, and small intestine submucosal tissues (SIS) were decellularized using previously established mechanical and enzymatic methods (21,22). For preparation of lung and liver dECM, the tissue was diced (∼0.5 mm pieces) and placed in a hypotonic solution (10 mM Tris and 5 mM EDTA) followed by a hypertonic solution (50 mM Tris, 1 M NaCl, and 10 mM EDTA). The tissue was then enzymatically digested using a 0.02% trypsin with 0.05% EDTA solution for 2 hours at 37ºC. The tissue was then placed in 4% sodium deoxycholate (SDC) to further solubilize and remove cell components. After thoroughly washing the tissue twice to remove residual SDC, tissues were placed in 3% Triton X-100. Tissues were then thoroughly washed and placed in a solution containing 0.1% peracetic acid with 4% ethanol. Once the tissue had undergone a final washing in PBS and water to completely remove any residual detergents and acid, it was then frozen, lyophilized and cryomilled to obtain a powder for use in hydrogels. For preparation of SIS dECM, gastrointestinal fluids were flushed from porcine small intestines and cut open lengthwise. The mucosal and muscular layers were then removed by mechanically delaminating the tissue. The submucosa was cut into 6 in. pieces and further decellularized and sterilized in a solution of 4% ethanol and 0.1% PAA. The decellularized tissue was then washed using 1x PBS and Type 1 water, after which it was lyophilized and cryomilled in preparation for hydrogel formulation. To prepare dECM hydrogels from the cryomilled dECM, the dECM powder was first digested. To do this, a 1 mg/mL pepsin solution in 0.01 N hydrochloric acid (HCl) was made. Cryomilled dECM was then digested in the pepsin solution at a concentration of 10 mg/mL for 48 hours on a magnetic stir plate. The hydrogels were prepared using a dECM concentration of 6 mg/mL. Digested dECM was then added to 1X PBS after which it was diluted at a 1:9 volume ratio of 10X PBS to dECM digest, followed by a dilution at a 1:10 volume ratio of 0.1 N NaOH to dECM digest, mixing between each addition. For example, for 250 uL of a 6 mg/mL hydrogel, 150 µL of the dECM digest was added to 68.33 µL of 1X PBS. Then, 16.67 µL of 10X PBS was added, followed by 15 µL of 0.1 N NaOH. The pre-gel was then incubated at 37^°^C for 30 minutes to achieve a viscoelastic hydrogel.

### Fluorescent labeling of AAV viral vectors

To visualize individual AAV viral vectors within these dECM hydrogels, AAV2, AAV6, and AAV8 viral vectors were fluorescently labeled with the amine-reactive fluorescent dye, Alexa Fluor Cy3 carboxylic acid succinimidyl ester (AF-Cy3) with excitation/emission (Ex/Em) at 555/572. As noted, GMP grade AAV was used throughout the described studies. Our lab has previously used this protocol to fluorescently label AAV viral vectors to measure their diffusion in human mucus (23,24). Before beginning the labeling protocol, stock viral vectors were diluted with 5% glycerol in PBS to prevent vector aggregation. A reactant mixture is then prepared by diluting 50 µL of the AAV stock in a solution containing 20 µL of PBS and 15 µL of 600 nM borate buffer (pH 8.3). A dye solution was prepared with 100 µg of AF555 stock dye reconstituted in 10 µL of DMSO. A 1.08 µL volume of this dye solution, along with 3.92 µL of DMSO was added to the reactant mixture. This mixture was constantly stirred and mixed in an end-over-end rotator at 4^°^C in the dark to allow the fluorescent dye molecules to adhere to the viral vector capsids. After 2 hours, the unreacted dye molecules are removed by buffer exchange into PBS using gel filtration chromatography via Sephadex gel columns. These columns were formed by mixing 1 g of commercial-grade Sephadex powder in 10 mL of PBS and allowing the solution to swell overnight. Microcolumns were filled with 400 uL of this solution, with the viral solution deposited on top and spun at 1000xg for 2 minutes. This process was repeated until the remaining eluate was a clear solution, indicating the absence of unreacted fluorescent dye molecules. The viral solution was then aliquoted into 10 µL aliquots for future use and stored at −80^°^ C.

### Preparation of PEGylated nanoparticles

Polystyrene nanoparticles were prepared in accordance with an established protocol routinely used by our group (22,25–27). Briefly, carboxylate modified fluorescent polystyrene spheres with a diameter of 20 nanometers were coated with polyethylene glycol at a high surface density using 5 kDa methoxy PEG-amine. Borate buffer (pH 8.3), N-Hydroxysulfosuccinimide sodium salt, and 1—ethyl-3-(3-dimethylaminopropyl) carbodiimide hydrochloride were used to activate and link the PEG-amine to the nanoparticles. After a series of mixing and washing steps, the particle size and zeta potential was measured using dynamic light scattering to confirm successful PEGylation.

### Multiple Particle Tracking Analysis

Multiple particle tracking was employed to evaluate the diffusion of fluorescently labeled AAV viral vectors in dECM hydrogels. To do this, we added 25 µL of the dECM hydrogel to be examined (liver, lung, or small intestinal submucosa) to a microscopy chamber and allowed it to gel at 37^°^ C for 30 minutes. After gelation, we added 1 µL of the fluorescently labeled AAV viral vector stock to be examined (AAV2, AAV6, or AAV8), along with 1 µL of 0.013% w/v PEGylated 20 nm nanoparticles as a size control. We then allowed the gels to equilibrate at room temperature for 30 minutes prior to imaging. The diffusion of AAV and the 20 nm PEGylated nanoparticles were then imaged in real-time using a Zeiss 800 LSM microscope with a 63x water-immersion objective and Zeiss Axiocam 702 camera at a frame rate of 33.33 Hz for 10 seconds. AAV and nanoparticle trajectories and diffusion rate were determined using a MATLAB-based image analysis software to track NP position over time. The time-averaged mean squared displacement (MSD(*τ*)) as a function of lag time, *τ*, was calculated from these trajectories as MSD(*τ*)=⟨[x(t+*τ*)−x(t)]^2^+[*y*(*t*+*τ*)−*y*(*t*)]^2^⟩. A representative measure of the diffusion rate was taken as the logarithm (base 10) of the MSD at a time scale of τ = 1 s (log_10_[MSD_1s_]). We performed each dECM hydrogel tracking experiments in triplicate with videos acquired in 5 distinct regions.

### A549 cell culture and AAV transduction

A549 cells acquired from ATCC were grown in with F-12K growth media supplemented with 10% v/v fetal bovine serum and 1% v/v penicillin-streptomycin seeded in multi-well plates. For all transduction studies conducted in 2D, AAV2-eGFP, AAV6-eGFP, and AAV8-eGFP were incubated for 2 hours at 37ºC in complete media prior to being introduced to the cells. Each AAV serotype was tested in individual wells in triplicate and additional control wells were reserved to set the baseline for transduction efficiency and cell viability. The multiplicity of infection (MOI), defined as the ratio of AAV viral particles (vp) relative to the number of cells, was set as 7.4 × 10^4^ vp/cell for each experiment and used to determine the dose for AAV with the following equation: number of cells per well x MOI = vp per well. From this, one can calculate the volume of AAV to be added to each well based on viral titer (in vp/mL) via the equation: (vp/well) / (vp/mL) = volume of AAV treatment. For the dECM incorporated 2D in vitro model, dECM hydrogels were seeded as a layer onto the surface of the 2D culture. Once A549 cells were fully enveloped in ECM, AAV-eGFP vectors are added apically and as such, AAV are required to diffuse through the hydrogel, simulating the physiological barrier of the ECM. Following a 48-hour incubation period to allow for viral transduction, fluorescent images were obtained via wide field and confocal microscopy to assess fluorescent signal distribution and intensity within the 2D in vitro model. GFP expression was quantified by image analysis via FIJI image software (ImageJ) using the automated thresholding function. Mean fluorescent intensity and number of successfully transduced cells were also evaluated using flow cytometry (BD FACS Celesta). Cells were prepped for flow cytometry first by being washed 2x with PBS. Cells were then incubated with 0.25% trypsin-EDTA to detach them from the plate and re-suspended in F-12K media prior to being centrifuged at 900xg for 2 minutes. The cells were then resuspended in PBS and stained with NIR Zombie dye (live-dead assay) which has excitation wavelength of 633 nm and an emission maximum of 746 nm (APC/Cy7) and incubated for 30 minutes on ice followed by another PBS wash step. The cells were then fixed in a 50% v/v paraformaldehyde solution in PBS for 30 minutes at a temperature of 4ºC and washed once more with PBS prior to being stored at 4ºC.

### A549 tissue spheroid culture and AAV transduction

A549 cells were grown in 200 µL of F-12K growth media supplemented with 10% v/v fetal bovine serum and 1% v/v penicillin-streptomycin seeded in ultra-low attachment 96-well round bottom plates at a density of 10,000 cells per well. The cells were grown in these plates for 7 days until they formed uniform cellular spheroids. AAV2-eGFP, AAV6-eGFP, and AAV8-eGFP vectors (MOI of 7.4 × 10^4^) were incubated for 2 hours at 37ºC in complete media prior to being introduced to the cells. For the dECM incorporated 3D in vitro model, dECM hydrogels were loaded into the ultra-low attachment 96-well round bottom plates, and fully formed spheroids were embedded within them. To introduce AAV to these cultures, 100 µL of the 200 µL of media surrounding the spheroids was removed and replaced with 100 µL of media containing AAV viral vectors at the specified MOI. The spheroids were incubated with the AAV for 48 hours to allow for transduction. After 48 hours, the spheroids were removed from the well and placed into serum-free media containing 0.25 mg/mL liberase TL and incubated for 20 minutes at 37^°^ C to break apart the spheroids. They were then centrifuged and the single cell suspension was washed to remove any residual AAV and liberase TL. The cells were then resuspended in a PBS wash and stained with NIR Zombie dye (live-dead assay) and incubated for 30 minutes on ice followed by another PBS wash step. The cells were then fixed in a 50% v/v paraformaldehyde solution in PBS for 30 minutes at a temperature of 4ºC and washed once more with PBS prior to being stored at 4ºC. Spheroids that had no AAV viral vectors introduced to them served as a control. Confocal microscopy was used to obtain images at the center of the spheroid and z-stack images to examine the GFP expression of the cells that indicated successful transduction of the cells.

### Statistics

The data collected was analyzed statistically using the GraphPad Prism Software. Data sets between groups were statistically analyzed with one-way analysis of variance (ANOVA) and a Tukey post hoc correction. All graphs display the median values and the 95^th^ down to the 5^th^ percentiles of data, with outliers included as well. Differences were considered statistically significant at a p value that was less than .05 (p < .05).

## Results

### Serotype-dependence of AAV diffusion through decellularized ECM

We first characterized the diffusion of AAV2, AAV6, and AAV8 within a model of the ECM. To capture the tissue-dependent chemical properties of the ECM, we employed decellularized ECM hydrogels as established in our previous work (21,22). Porcine lung, liver, and small intestine submucosal tissues (SIS) were collected and decellularized via chemical and mechanical means and brought to physiologically relevant pH levels to form ECM-based hydrogels. Fluorescently labeled AAV2, AAV6, AAV8, and 20 nm PEGylated nanoparticles were then added to measure their diffusion within hydrogels derived from each tissue type (**Fig. 1**). Previous work has established AAV labeled with this procedure retain their biological activity (28). PEGylated nanoparticles with ∼20 nm diameter were used as control for comparison to AAV viral particles similar in size (∼25 diameter) (29). Each dECM hydrogel type was seeded in custom imaging chambers, and fluorescently labeled AAV2, AAV6, and AAV8 viral vectors were seeded into separate wells for observation. PEGylated NPs were added to each well as a control, and diffusion rates for all AAV vectors and the NPs were obtained in the same regions of interest via fluorescence video microscopy.

**Figure 1.**
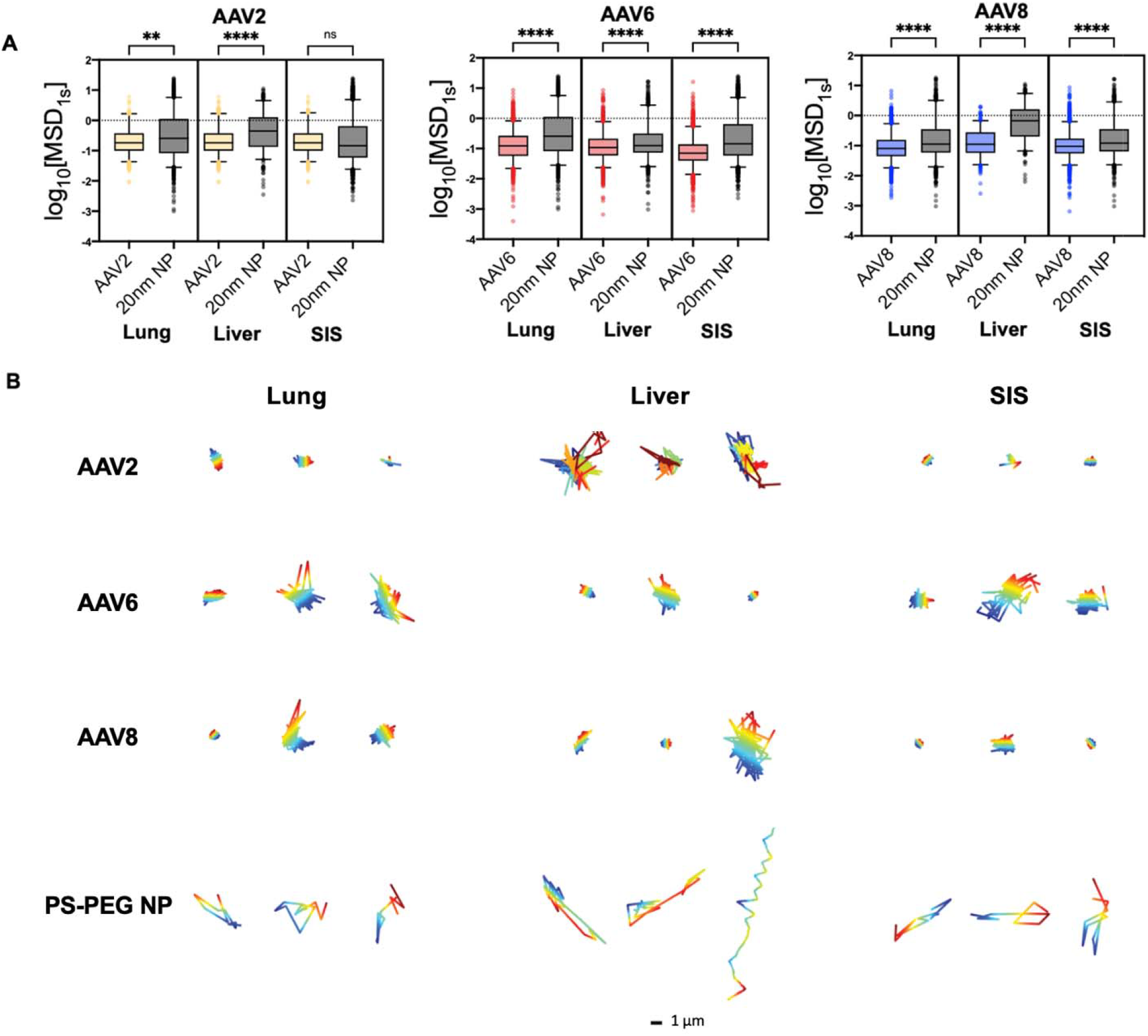
AAV diffusion through decellularized ECM hydrogels. (**A**) Diffusion of AAV2 (yellow), AAV6 (red), AAV8 (blue), and 20 nanometer NPs (black) represented as the median log based 10 of MSD at τ=1 s (log_10_[MSD_1s_]) in lung dECM, liver dECM, and SIS dECM hydrogels are shown (n=3 replicates). Horizontal bars represent median values. (**B**) Representative trajectories for all AAV vector serotypes and 20 nanometer NPs in all dECM hydrogel types. Traces show 10 s of motion with a color scale to indicate time. The scale bar = 1μm. Data sets in (**A**) statistically analyzed with Kruskas-Wallis test: **** p□<□0.0001. At least 5 videos in different regions of the gel were acquired with ≥150 particles tracked for each experimental condition.

Each video was analyzed via multiple particle tracking (MPT) analysis yielding individual particle trajectories. From these trajectories, the mean squared displacement (MSD) was determined, which is representative of the rate of diffusion of AAV and nanoparticles within the ECM. Based on these values, the diffusion rates of AAV6 and AAV8 were significantly lower than the similarly sized PEGylated NPs in all dECM types **(Fig. 1A)**. AAV2 displayed a similar trend in both the lung and liver dECM hydrogels but did not display a significant difference in diffusion rate in SIS dECM hydrogels **(Fig. 1A)**. These diffusion rates indicate a slower and more hindered movement of AAV when compared to a neutrally charged densely PEGylated nanoparticle and the reduced transport of AAV can likely be attributed, at least in part, to binding to the ECM. To visualize these effects, 3 representative diffusion trajectories taken from videos collected from each dataset were produced **(Fig. 1B)**. Across all ECM types and AAV serotypes tested, these trajectories display a far more localized and restricted trajectory pattern for AAV in comparison to the PEGylated NP control across. Additionally, the degree to which each AAV is trapped within the dECM varies significantly depending on the tissue of origin. For example, AAV2 particle mobility appears the least hindered in liver-derived dECM compared to other tissue types.

In **Fig 2**, we further compared AAV diffusion across the different serotypes and dECM hydrogels tested. We note these data are from the same experiments shown in **Fig 1** but re-plotted to allow for comparisons between serotypes in dECM gels from distinct tissues. We found the diffusion rate of AAV2 was the highest in lung and liver dECM, subsequently followed by AAV6, and then AAV8, having the lowest rate amongst the three serotypes **(Fig. 2A)**. However, in SIS this trend varied slightly, with AAV2 having the highest diffusion rate followed by AAV8, and AAV6 having the lowest rate amongst the three serotypes **(Fig. 2A)**. These findings suggest the mobility of AAV in the ECM is both dependent on the specific AAV serotype and tissue of origin for the ECM barrier. Further, we evaluated the diffusion of AAV6 in collagen to compare to its mobility to that found in dECM gels (**Fig. 2B**). Interestingly, we find AAV6 achieves significantly greater diffusion in collagen gels as compared to dECM which suggests the non-collagen components (e.g. laminin, fibronection, GAGs) may facilitate AAV trapping.

**Figure 2.**
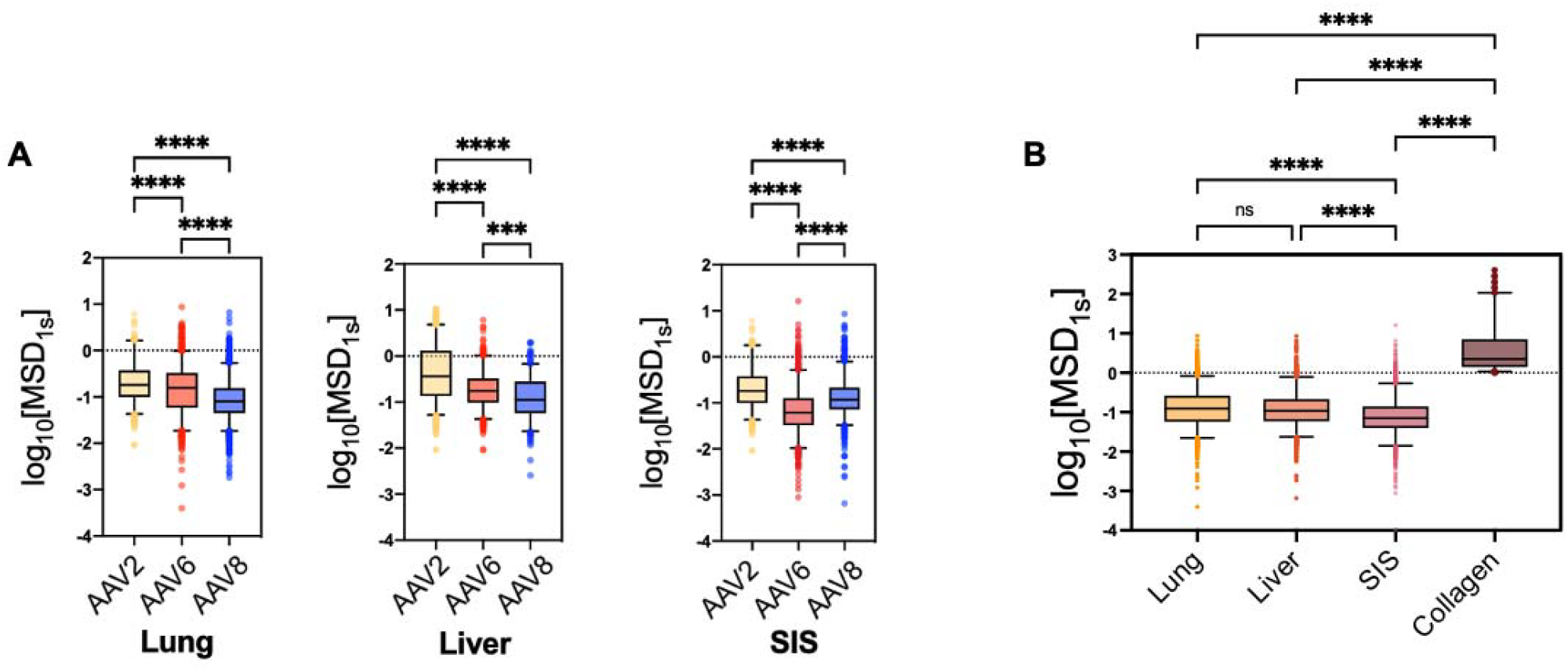
Serotype- and tissue-dependence of AAV diffusion through the ECM. (**A**) Virus diffusion as represented by log based 10 of MSD at τ=1 s (log_10_[MSD_1s_]) of AAV2 (yellow), AAV6 (red), AAV8 (blue) in lung, liver, and SIS dECM hydrogels. (**B**) Diffusion of AAV6 as measured by log_10_[MSD_1s_] in collagen, lung dECM, liver dECM, and SIS dECM hydrogels. Data sets statistically analyzed with Kruskas-Wallis test: **** p□<□0.0001, ns = not significant. At least 5 videos in different regions of the gel were acquired and analyzed in each experiment.

### AAV transduction in cells overlayed with decellularized ECM

To further test the functional impact of the ECM on AAV and its ability to deliver genetic cargo, we established a dECM incorporated 2D *in vitro* model. To accomplish this, we deposited a layer of dECM hydrogel onto the surface of A549 cells grown in 2D. In this format, dECM hydrogel coat the surface of the cells and form a barrier which the AAV viral vectors must pass through prior to transduction **(Fig. 3A)**. For these studies, lung dECM hydrogels were used due to the A549 cells being a lung-derived cell line. AAV6-eGFP viral vectors were introduced at MOI of 7.4 × 10^4^ vp/cell. After 48 hours, the hydrogels were chemically degraded and the cells were collected, stained and fixed for analysis via flow cytometry.

**Figure 3.**
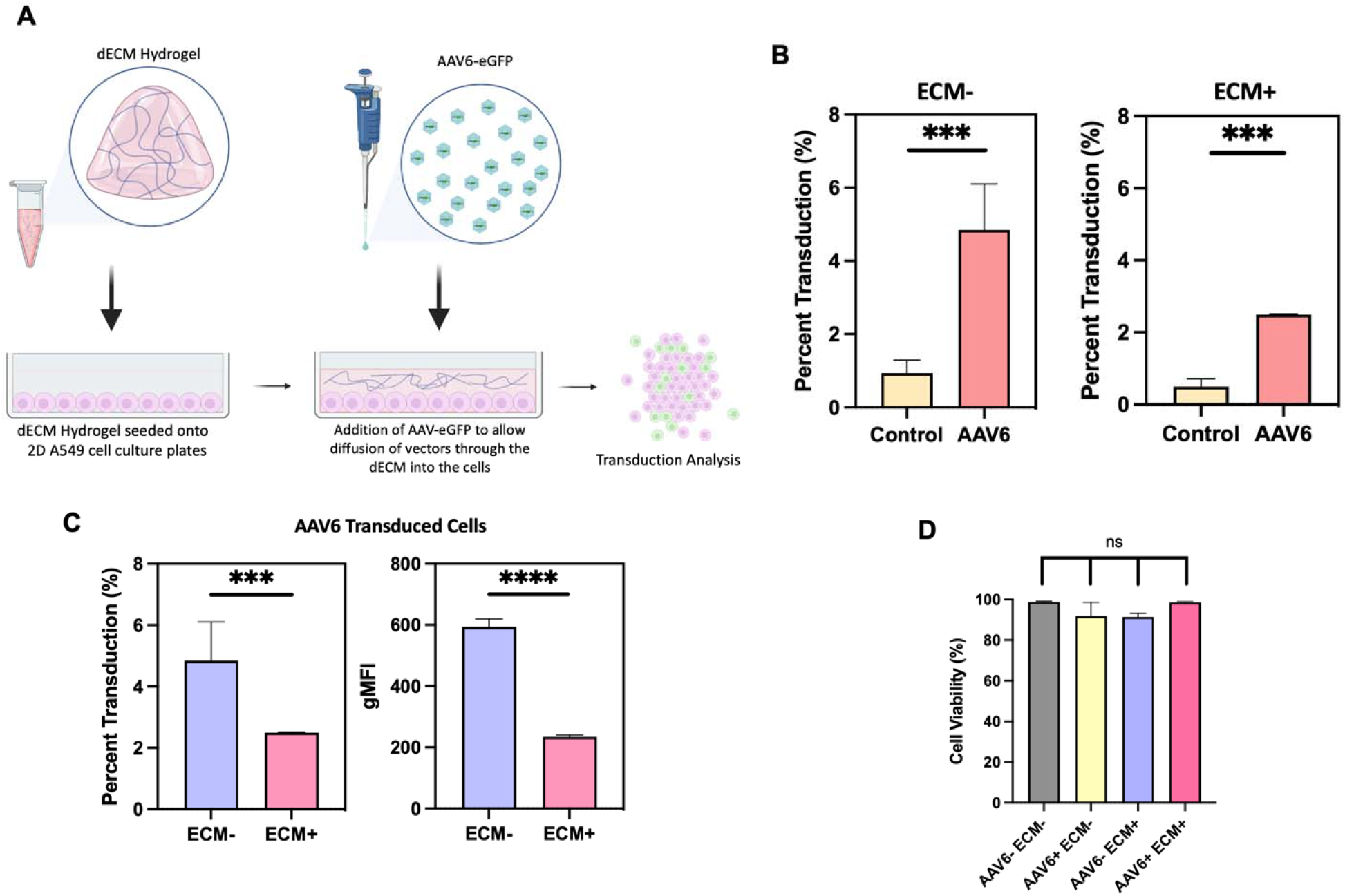
AAV transduction in cells cultured in 2D and overlayed with dECM hydrogels. (**A**) Schematic of 2D cell culture immersed in a layer of the lung dECM hydrogel. **(B)** AAV6 transduction of 2D A549 cell culture without ECM (ECM-) and in cultures immersed in a hydrogel composed of 6mg/mL lung dECM (ECM+). AAV6 mediated eGFP expression were evaluated via flow cytometry. AAV6-treated groups had a sample size of n=3 with a cell count ∼50,000 cells each. Untreated (control) groups had a sample size of n=2 with a cell count of ∼100,000 cells each. **(C)** Comparison of AAV6 transduction with and without the presence of the 6mg/mL lung dECM hydrogel barrier. Geometric mean fluorescence intensity from viable cells collected in A549 cultures both immersed and devoid of the 6mg/mL lung hydrogels obtained via flow cytometry. **(D)** Cell viability values obtained via cell staining with Zombie NIR stain and flow cytometry analysis. Data sets in (**B,C**) analyzed via unpaired t-test with Welch’s correction: \*\*\**p* < 0.001. Data in (**D**) statistically analyzed with one-way analysis of variance (ANOVA) and a Tuke post hoc correction: \*\*\**p* < 0.001, \*\*\*\**p* < 0.0001.

Quantification of percent transduction revealed that there was a significant increase in GFP expression in cells treated with AAV6-eGFP **(Fig. 3B)**. When comparing percent transduction between AAV6-eGFP vectors that had to travel through the dECM hydrogel and those that did not, we see that the presence of the dECM hydrogel significantly decreased the percentage of transduced cells and resulted in lower geometric mean fluorescence intensity (gMFI) **(Fig. 3C)**. There were also no significant differences seen in A549 cell viability when cultured in the presence of an ECM barrier **(Fig. 3D)**.

### AAV transduction in 3D tissue spheroids

We next wanted to evaluate the transduction profile for each AAV serotype within a cellular spheroid model. Given their 3D architecture and endogenous expression of ECM components (30–32), we hypothesized tissue spheroids would present a physicochemical barrier to AAV diffusion and transport to target cells. To test this, we seeded the A549 cell line in ultra-low attachment 96-well round bottom plates to initiate growth of 3D cellular spheroids. A549 tissue spheroids were then subsequently treated with AAV2-eGFP, AAV6-eGFP, and AAV8-eGFP (MOI of 7.5 × 10^4^, 48-hour incubation) and evaluated via fluorescence microscopy. GFP fluorescence was observed in spheroids transduced with each serotype in comparison to untreated controls **(Fig. 4A)**. No significant difference was observed in percent transduction or overall fluorescent intensity for all AAV serotypes **(Fig. 4B, 4C)**. These results establish that each AAV serotype can overcome the intra- and extracellular barriers to gene delivery presented in this model.

**Figure 4.**
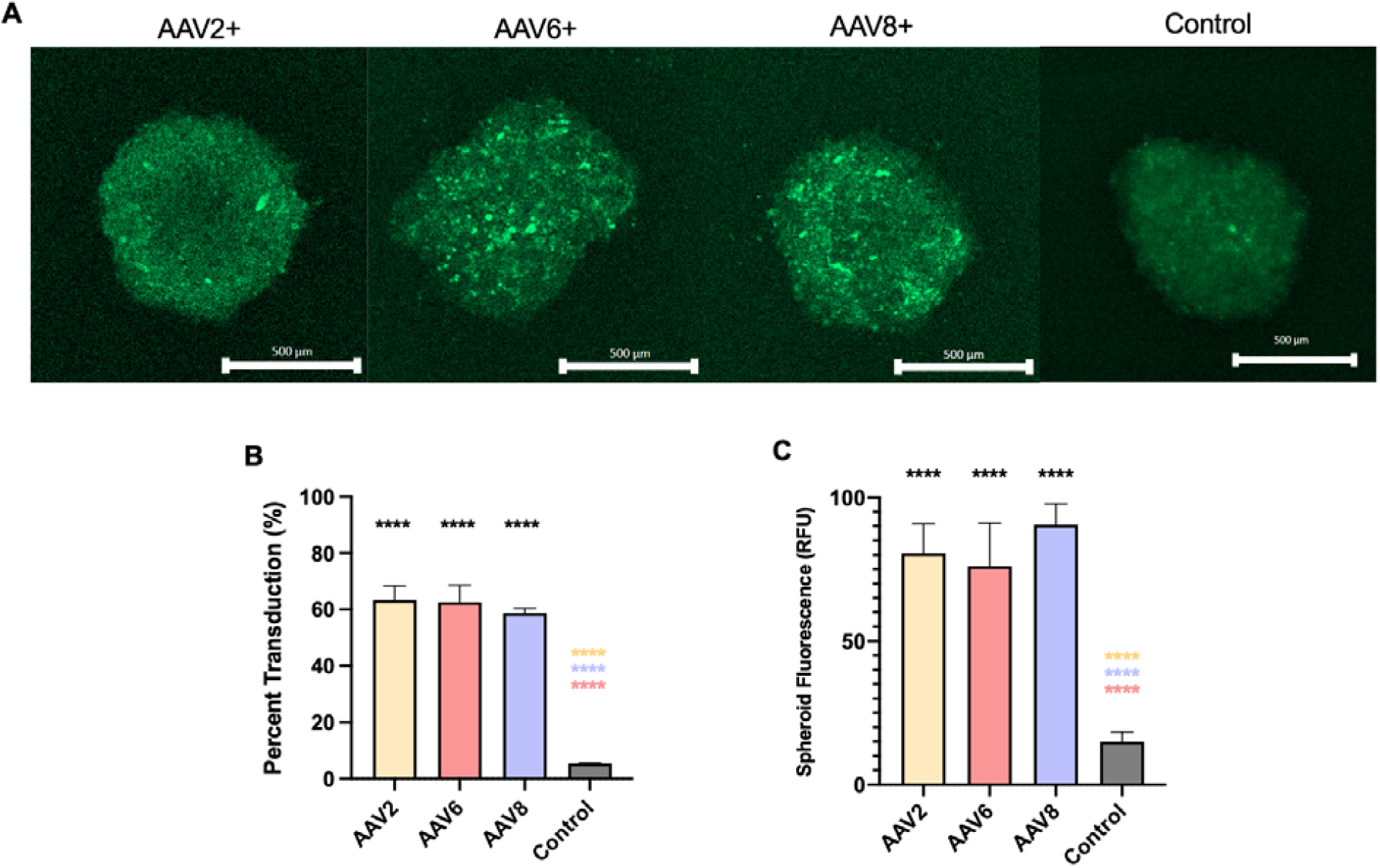
AAV transduction in 3D Tissue Spheroids. (**A**) In vitro fluorescent confocal images of AAV transduction in A549 cells in 3D cell culture. Images displayed are representative for A549 tissue spheroids transduced with AAV2, AAV6, and AAV8 viral vectors containing the eGFP transgene. Transduction studies conducted in 3D tissue spheroids comprised of A549 cells that were grown for a period of 7 days and transduced with vectors at a multiplicity of infection of 75,000 vp per cell. Fluorescent images were taken 48 hours post transduction using confocal microscopy (10X, dry immersion, Scale Bar = 100 µm). (**B**) Percent transduction determined via ImageJ by comparing area of GFP+ and DAPI+ regions from fluorescence imaging of A549 tissue spheroids (n=3 replicates). (**C**) Fluorescent images were analyzed via ImageJ to obtain measurements of spheroid fluorescence (values measured in relative fluorescent units). Data set statistically analyzed with one-way analysis of variance (ANOVA) and a Tukey post hoc correction: \*\*\*\**p* < 0.0001.

### Decellularized ECM negatively impacts AAV transduction in 3D tissue spheroids

We next established a dECM incorporated 3D *in vitro* model for further testing on the functional role of ECM barrier function in AAV transduction efficiency. Briefly, we established a A549 *in vitro* spheroid model as used previously (**Fig 4**). We then deposited fully formed cellular spheroids into dECM hydrogels, allowing the hydrogel to envelop all surfaces of the spheroid **(Fig. 5A)**. This experimental format simulates a stromal ECM barrier surrounding target cells. Like before, AAV6 (MOI of 7.4 × 10^4^) were introduced into the model and 48 hours were allowed for transduction. After 48 hours, the hydrogels were chemically degraded and the spheroids were broken up into single cells and collected for analysis via flow cytometry. When comparing percent transduction of the spheroids between AAV6-eGFP vectors that had to travel through the dECM hydrogel and those that did not, we see that the presence of the dECM hydrogel significantly decreased the percent of transduced cells and geometric mean fluorescence intensity (gMFI) **(Fig. 5B)**. We do note a decrease in cell viability was seen in spheroids that were enveloped in the dECM hydrogel (**Fig. 5C)**.

**Figure 5.**
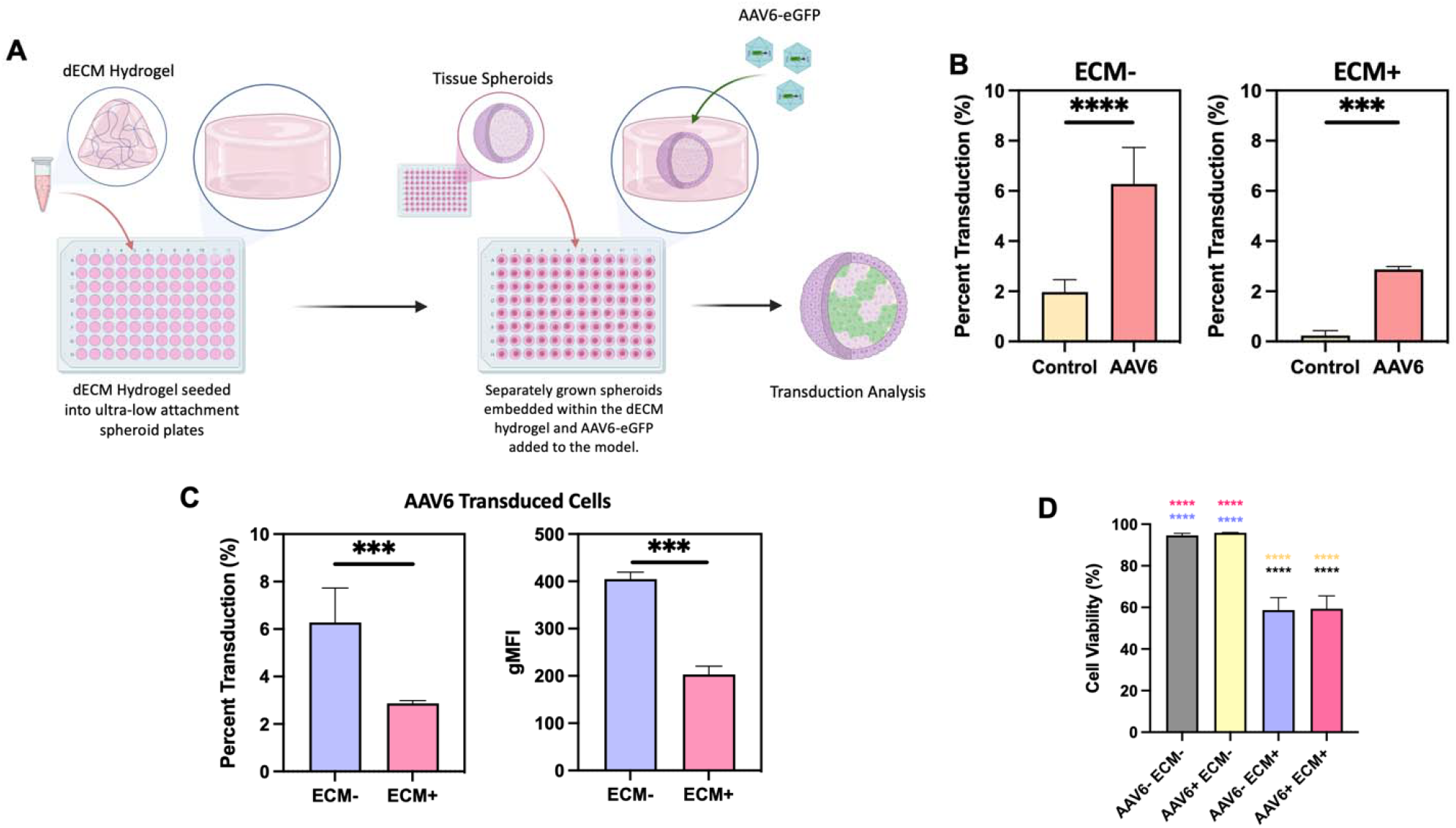
AAV transduction in dECM embedded 3D tissue spheroids. (**A**) Schematic of 3D tissue spheroids embedded in a layer of the lung dECM hydrogel. **(B)** AAV6 transduction of 3D A549 tissue spheroids in media (ECM-) and spheroids embedded in a hydrogel composed of 6mg/mL lung dECM (ECM+). AAV6 mediated eGFP expression was evaluated via flow cytometry. Percent transduction shown for viable cells transduced with and without the presence of ECM as well as untreated controls. All groups were repeated in triplicate with a cell count of ∼15,000 per condition. **(C)** Geometric mean fluorescence intensity of GFP expression for in AAV6 treated A549 cultures both immersed and devoid of the 6mg/mL lung hydrogels obtained via flow cytometry and compared. **(D)** Cell viability evaluated via Zombie NIR staining and flow cytometry analysis. Data sets in (**B,C**) analyzed via unpaired t-test with Welch’s correction: \*\*\**p* < 0.001. Data in (**D**) statistically analyzed with one-way analysis of variance (ANOVA) and a Tukey post hoc correction: \*\*\**p* < 0.001, \*\*\*\**p* < 0.0001.

## Discussion

In these studies, we evaluated the effect of the ECM as a barrier to AAV gene delivery. Defining the effect of ECM on AAV transport within tissues and transduction in target cells has important implications in the design of AAV-based *in vivo* gene therapies. We first established the diffusion profiles of three serotypes (AAV2, AAV6, AAV8) in a decellularized ECM hydrogel model from three distinct tissues (lung, liver, SIS). It is important to note that dECM hydrogels do not replicate *in vivo* tissue architecture. Rather, the dECM model allows us to systematically compare how differences in ECM composition influence barrier function towards AAV. Given the size of AAV (∼25 nm) compared to interstitial spacings in the ECM, we suspect reduced transport of AAV particles through the ECM will be primarily dictated by specific and non-specific interactions with ECM associated components. We hypothesized that differences in ECM and serotype behavior (e.g. binding to ECM proteins and/or proteoglycans) would significantly impact AAV penetration through the ECM barrier. Moreover, comparisons of AAV6 diffusion in dECM and collagen suggest non-collagen ECM components (e.g. laminin, fibronectin, GAGs) significantly influence the barrier function of ECM towards AAV.

We observed that the diffusion rates of all serotypes were consistently lower than a virus-sized 20 nm nanoparticle in all tissue types, except for 1 case for AAV2 in SIS dECM hydrogel. The comparison to a neutrally charged and similarly sized nanoparticle control indicates that hindered AAV diffusion through the ECM microenvironment is most likely due to binding effects between the viral capsids and polymer fibers within the ECM. We also found that the diffusion of each AAV serotype differed across the various dECM hydrogel tissue types with diffusion rate consistently the highest for AAV2. This supports the idea that serotype selection may play a role in determining AAV transport through and distribution within target tissues. Relative to other serotypes, the improvements in mobility of AAV2 also suggests binding to heparan sulfate-terminated glycosaminoglycans (GAGs) may not pose a substantial barrier to transport for AAV2. Conversely, AAV8 had the least mobility compared to AAV2 and AAV6 in dECM hydrogels which is notable given its lack of reported affinity for sulfated and/or sialylated GAGs. However, it is possible other non-specific interactions with ECM components, such as collagen, fibronectin, and laminin, may have slowed the transport of AAV8. We note as well the LamR receptor acquired from the cells within each tissue could be present in dECM via association to laminins leading to the observed trapping of AAV8. Overall, these studies suggest adsorption and binding of AAVs to ECM could be modulated by serotype-dependent properties of the AAV capsid.

Incorporation of dECM hydrogels in a 2D and 3D *in vitro* model enabled further investigation into the functional impact of the ECM barrier on AAV transduction. Notably, tissue spheroids were readily transduced by all 3 serotypes in the absence of an outer layer of dECM suggesting endogenous expression of ECM and/or intracellular junctions were not a limiting factor in AAV transduction. Encasing these spheroids within our dECM hydrogels lead to ≥50% reductions in percentage of transduced cells and fluorescence intensity within individual cells. Thus, we reason the presence of a stromal ECM barrier significantly hindered the ability of AAV particles to reach the embedded spheroids leading to an apparent reduced transduction efficiency. We note the procedure of embedding of the spheroids in dECM, which required manual handling and placement in a well containing dECM precursors, lead to a reduction in cell viability. However, we have limited comparisons of transduction efficiency for the viable cell population in the dECM-free and dECM-embedded spheroids. We also recognize dECM-embedded cells may phenotypically differ from cells grown in dECM-free conditions, due to cell-ECM interactions (33–35), which may influence receptor-mediated endocytosis and AAV transgene expression.

## Conclusions

This study demonstrates that the extracellular matrix (ECM) microenvironment significantly impedes AAV diffusion and transduction in vitro. These findings underscore the importance of incorporating ECM components into *in vitro* models used to assess AAV gene vectors. AAV gene vectors are in clinical development for several indications, such as liver fibrosis and neurodegenerative disorders, requiring delivery in organs enriched in ECM posing a significant challenge in achieving widespread gene delivery to disease-affected cells. These results pave the way for research aimed at overcoming ECM-associated barriers to enhance the efficiency of AAV-based gene therapies.

## Acknowledgements

We acknowledge the BioWorkshop core facility in the Fischell Department of Bioengineering at the University of Maryland for use of their flow cytometer. The contributions of the NIH authors were made as part of their official duties as NIH federal employees, are in compliance with agency policy requirements, and are considered Works of the United States Government. However, the findings and conclusions presented in this paper are those of the author(s) and do not necessarily reflect the views of the NIH or the U.S. Department of Health and Human Services, nor does mention of trade names, commercial products, or organizations imply endorsement by the U.S. Government.

