## Supplemental Information for "Barrier Function of the Extracellular Matrix in AAV Gene Therapy"

**Supporting Information**

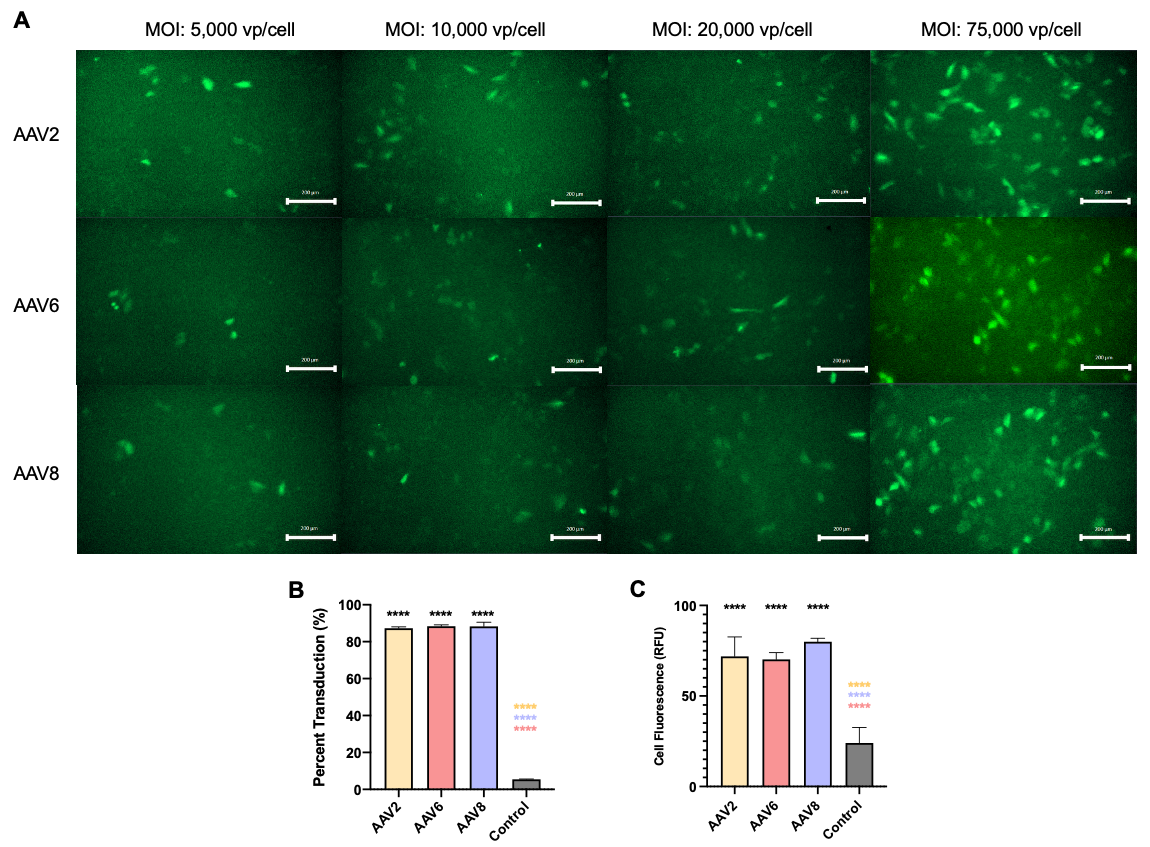

***Figure S1*** | **AAV transduction in 2D A549 cell culture.**

**(A) I**n vitro fluorescent images of AAV transduction in A549 cells in 2D cell culture. Images displayed are representative for AAV2, AAV6, and AAV8 viral vectors containing the eGFP transgene. Transduction studies conducted at 60% cell confluence with an MOI of 5,000 viral particles (vp) per cell, 10,000 vp per cell, 20,000 vp per cell and 75,000 vp per cell. Images were taken 48 hours post transduction using wide field microscopy (10X, dry immersion, Scale Bar = 100 µm). **(B)** Percent transduction of 2D cell culture values obtained by fluorescent image analysis in comparison to DAPI stained images of the A549 cultures. **(C)** Fluorescent images then were analyzed via ImageJ to obtain measurements of cell fluorescence (values measured in relative fluorescent units).
